# Synergistic Behavioral and Neuroplastic Effects of Psilocybin-NMDAR Modulator Administration

**DOI:** 10.1101/2024.11.28.625811

**Authors:** Tom Ben-Tal, Ilana Pogodin, Alexander Botvinnik, Tzuri Lifschytz, Uriel Heresco-Levy, Bernard Lerer

## Abstract

The full therapeutic potential of serotonergic psychedelics (SP) in treating neuropsychiatric disorders, such as depression and schizophrenia, is limited by possible adverse effects, including perceptual disturbances and psychosis, which require administration in controlled clinical environments. This study investigates the synergistic benefits of combining psilocybin (PSIL) with N-methyl-D-aspartate receptor (NMDAR) modulators D-serine (DSER) and D-cycloserine (DCS) to enhance both efficacy and safety. Using ICR male mice, we examined head twitch response (HTR), MK-801-induced hyperlocomotion, and neuroplasticity related synaptic protein levels in the frontal cortex, hippocampus, amygdala, and striatum. Our results indicate that PSIL significantly increased HTR—a surrogate measure for hallucinogenic effects—which was reduced by the co-administration of DSER or DCS in a dose-dependent manner. Similarly, combining PSIL with DSER or DCS significantly decreased MK-801-induced hyperactivity, modeling antipsychotic effects. Neuroplasticity-related synaptic protein assays demonstrated that the PSIL-DSER combination enhanced GAP43 expression over all 4 brain examined and overall expression of the 4 assayed synaptic proteins in the hippocampus, while PSIL-DCS elevated PSD95 levels across all 4 brain regions, suggesting a synaptogenic synergy. These findings support the hypothesis that combinations of SP with NMDAR modulators could optimize the therapeutic potential of SP by mitigating adverse effects and enhancing neuroplasticity. Future studies should focus on refining administration protocols and evaluating translational applicability for broader clinical use.

## 1. Introduction

The use of classical serotonergic psychedelics (SP) is currently advocated for an increasing array of neuropsychiatric disorders, including depression, anxiety, post-traumatic stress disorder, addictions ^1, 2^ and, more recently, schizophrenia ^3^. This potentially wide therapeutic range can be explained by the capacity of SP to induce rapid, long-term changes in neuronal plasticity involving increased levels of neurotrophic proteins and promotion of dendritic spine growth and synapse formation ^4–7^.

Nevertheless, it is presumed that SP efficacy requires psychedelic effects, including alterations in consciousness and sensory perception, although this view is not held by all authors ^8, 9^. Moreover, although generally well tolerated in clinical and research settings, reported SP-induced serious adverse events include exacerbation of depression, suicidal behavior, and psychosis ^10^. These drawbacks presently mandate a clinical SP administration paradigm that includes monitoring, environmental manipulation, and costly psychological support in specialized inpatient settings ^1^, which impose significant barriers to widespread SP utilization.

We have previously hypothesized that concurrent administration of SP and N-methyl-D-aspartate receptor (NMDAR) modulators may synergistically upgrade SP efficacy and attenuate SP-induced unpleasant subjective reactions and side effects ^11^. NMDARs play a critical role in neuroplasticity, meta-plasticity, long-term potentiation (LTP), learning, and memory ^12^. Antipsychotic and pro-cognitive effects may be produced by modulators acting at the glycine modulatory site (GMS) on the NMDAR NR1 subunit ^13, 14^ while, paradoxically, antidepressant effects have been reported with both NMDAR agonists and antagonists ^15^. Moreover, although SP do not have direct affinity for glutamate (GLU) receptors, glutamatergic neurotransmission plays a significant role in their overall downstream effects. The activation by SP of postsynaptic serotonin 2 A receptors (5HT2AR) on pyramidal neurons leads to a GLU-dependent increase in the activity of neurons in the frontal cortex, subsequently modulating prefrontal network activity ^16, 17^. The resulting downstream signaling cascades involve ionotropic NMDAR and amino-3-hydroxy-5-methyl-isoxazolepropionic acid receptors (AMPAR) and increased AMPAR/NMDAR ratios ^18^. AMPAR activation may potentiate brain-derived neurotrophic factor-tropomyosin receptor kinase B (BDNF-TrkB) and mammalian target of rapamycin (mTOR) signaling, thus upregulating the expression of neuroplasticity-related genes and protein synthesis of synaptic components, ultimately leading to rapid and long-lasting synaptogenesis ^7, 17^.

Relevant to safety issues, 5HT2AR-NMDAR interactions are known to modulate neuronal excitability in the prefrontal cortex and have been proposed as a potential target for antipsychotic drugs ^19^.

NMDARs blockade upregulates cortical expression of 5HT2AR, which is implicated in psychosis. Genetic depletion of the NMDAR NR1 subunit and noncompetitive NMDAR antagonists potentiate the activation of serotoninergic receptors, upregulating 5HT2AR and leading to cortical hyperexcitability ^20, 21^. On the contrary, positive modulators of NMDAR could inhibit serotoninergic activation ^22^, suggesting an antipsychotic potential. Overall, these converging lines of evidence support the concept that combined SP-NMDAR modulator treatment may offer efficacy as well as safety advantages.

We propose to first focus research on the combined administration of psilocybin with the prototypical NMDAR-GMS modulators D-serine (DSER) and D-cycloserine (DCS). Accumulating data indicate that all these compounds have significant neuroplastic and clinical effects. Psilocin, the active metabolite of psilocybin, has been reported to induce changes in neuroplasticity, including neurogenesis, that were accompanied by extinction of conditioned fear-related behaviors ^6, 23, 24^.

Significant benefits in major depressive disorder for 1-2 psilocybin (∼25mg) administration sessions have been documented ^25, 26^. Nevertheless, psilocybin-induced alterations in perception and potential suicide risk ^27^ represent significant drawbacks.

DSER is an endogenous obligatory NMDAR co-agonist at GMS that does not bind to other known targets ^28^. An acute DSER dose has antidepressant effects in animal models of depression that are mediated by rapid AMPAR-induced mTOR signaling and increased BDNF levels ^29, 30^. DSER administration results in increased extra- and intracellular brain DSER levels and can enable LTP and/ or reverse age-related deficits in LTP expression and learning performance ^31, 32^. In mice, DSER administration increases hippocampal cell proliferation, the density of neural stem cells, and the survival of newborn neurons ^33^. A role as a cognitive enhancer (30) and in the treatment for schizophrenia negative symptoms ^34, 35^ has been suggested for DSER.

DCS is a *Streptomyces*-isolated broad-spectrum antibiotic approved for tuberculosis treatment, used since the 1950s, usually at 500-1000 g/day regimens, in millions of individuals. The discovery of a partial agonist role for DCS at GMS in the 1980s led to the characterization of its impact on the facilitation of cortical neuroplasticity ^36^. In brain injury animal models, DCS can reverse synaptic plasticity alterations via improved LTP, restored BDNF levels, and increased dendritic spine density ^37–39^. DCS enhances extinction of conditioned fear ^40, 41^ and shows efficacy in a variety of models related to deficient fear extinction and alcohol withdrawal ^42–45^. In humans, treatment with DCS administered in conjunction with cognitive behavioral therapy sessions enhances the efficacy of exposure therapy for various forms of maladaptive fear, including social anxiety, obsessive-compulsive disorder, and panic disorder ^46^. Recently, antidepressant and anti-suicidal effects were found with DCS in treatment-resistant major depressive disorder ^47, 48^ and bipolar depression ^49^.

To investigate our hypotheses, the present study assessed in experimental murine models the effects of concurrent acute administration of DSER or DCS and psilocybin on parameters relevant to SP treatment. The specific parameters examined were head twitch response (HTR), MK-801-induced hyperlocomotion, and synaptic protein levels. HTR is a rhythmic paroxysmal rotational head movement that occurs in rodents in response to 5-HT2AR activation ^50, 51^, correlates with the psychedelic trip in humans, and is a well-established animal behavioral assay for hallucinogenic-like effects ^52^. MK-801-induced hyperlocomotion, a putative model for acute antipsychotic effects, is used as a pharmacological model of schizophrenia, mimicking its positive symptoms ^53, 54^.

Furthermore, we evaluated post-treatment levels of the synaptic proteins GAP43, PSD95, synaptophysin, and SV2A in the frontal cortex, amygdala, hippocampus, and striatum. We hypothesized that administration of the combined treatments (psilocybin + DSER or psilocybin + DCS) would down-regulate HTR and MK-801-induced hyperlocomotion while increasing synaptic protein levels.

## 2. Methods

### a. Animals

The study was carried out on male ICR mice (30.0 ± 2.0 g). The animals were housed in a temperature-controlled environment (25 ± 2 °C, relative humidity 55 ± 15%) with a 12-h light/dark cycle. All mice were fed a standard laboratory diet of nutrient-rich pellets ad libitum. The mice were allowed to acclimatize for a week before starting the experiment. Separate sets of mice were used for the head twitch response (HTR), MK-801-induced hyperactivity, and synaptic protein experiments. The sample size was estimated based on prior studies. Since the tests performed were based on objective parameters, formal blinding was not applied. Experiments were approved by the Authority for Biological and Biomedical Models, Hebrew University of Jerusalem, Israel (Animal Care and Use Committee Approval Numbers: MD-22-17000-4 and MD-24-17494-4). All efforts were made to minimize animal suffering and the number of animals used.

### b. Drugs

Drugs were administered by intraperitoneal (i.p.) injection in a standard injection volume of 300 µl. Mice were divided into the following treatment groups: Vehicle (VEH): 0.9% saline; Psilocybin (PSIL): 4.4 mg/kg dissolved in the vehicle, D-seine (DSER) 3000 mg/kg or D-Cycloserine (DCS) 320 mg/kg dissolved in the VEH and combinations of PSIL and DSER or PSIL and DCS at the same concentrations. The experiments with DSER and DCS were conducted separately. Additional experiments were conducted with different doses of PSIL and DSER or DCS; the results are presented separately. PSIL was supplied by Usona Institute, Madison, WI, USA, was determined by AUC at 269.00 nm (UPLC) to contain 98.75% psilocybin and stored in a cool, light-sealed safe. DSER was supplied by Merck Sigma Aldrich, Rehovot, Israel, and was stored at room temperature of 37°C. DCS was supplied by Merck KGaA, Darmstadt, Germany and stored at -20°C. PSIL, DCS and DSER were dissolved in a 0.9% saline vehicle.

### c. Head-twitch response (HTR)

Assessment by electromagnetic generation of the rapid side-to-side head twitch that characterizes HTR requires the installation of small magnets in the outer ears of the mice as previously described ^55^. For this purpose, small neodymium magnets (N50, 3 mm diameter × 1 mm height, 50 mg), which were attached to the top surface of aluminum ear tags (supplied by Mario de la Fuente Revenga, PhD. of Virginia Commonwealth University), were used. The ear tags were placed through the pinna antihelix and laid in the interior of the antihelix, resting on top of the antitragus, leaving the ear canal unobstructed. This procedure was performed by simple restraint and immobilization of the mouse’s head. The ear tags were well-tolerated by the mice, and signs of ear tissue damage in the form of redness were rare. After a 7-day recovery period, the ear-tagged animals were placed inside a magnetometer apparatus (supplied by Mario de la Fuente Revenga, PhD., of Virginia Commonwealth University) immediately after IP injection of VEH, PSIL 4.4 mg/kg, DSER 3000 mg/kg DCS 320 mg/kg or a combination of PSIL and DSER or PSIL and DCS. Additional experiments were performed with a DSER dose of 1500 mg/kg (and PSIL 1.5 mg/kg) and DCS doses 22.2mg/kg, 176.19mg/kg, and 440.47 mg/kg (and PSIL 4.4 mg/kg). The output was amplified (Pyle PP444 phono amplifier) and recorded at 1000 Hz using a NI USB-6001 (National Instruments, US) data acquisition system.

Recordings were performed using a MATLAB driver (MathWorks, US, R2021a version, along with the NI myDAQ support package) with the corresponding National Instruments support package for further processing. A custom MATLAB script was used to record the processed signal, which was presented as graphs showing the change in current as peaks (mAh). A custom graphical user interface created in our laboratory was used to further process the recording into an Excel spreadsheet.

### d. MK-801-induced hyperactivity test

Mice were injected IP with VEH, PSIL 4.4 mg/kg, DSER 3000 mg/kg, or a combination of both or with VEH, PSIL 4.4 mg/kg, DCS 320 mg /kg, or a combination of both. After 30 minutes, the mice were injected IP with MK-801 0.5 mg/kg, and 30 minutes later, they were placed in the open field arena. The open field apparatus consisted of 4 square wooden arenas (50 × 50 × 40 cm) with white walls and a black floor. Mice were placed individually in the center of the open field and allowed to freely explore the apparatus for 60 min. A camera was used to monitor movement. The total distance traveled (centimeters) was measured by the Ethovision XT-17 Video Tracking System (Noldus Information Technology BV). After each test, the arena was cleaned with a 70% alcohol solution.

### e. Synaptic Protein Neuroplasticity Markers

The frontal cortex, hippocampus, amygdala, and striatum were dissected 12 days after IP injection of VEH, PSIL 4.4 mg/kg, DCS 320 mg/kg, or a combination of PSIL and DCS for analysis of levels of GAP43, PSD95, synaptophysin, and SV2A.

For the PSIL/DSER study mice were treated with VEH, PSIL (4.4 mg/kg), DSER (3000 mg/kg) or a combination of both on day 1. On day 8, a week after the 1st treatment, mice received the same treatments again. 11 days after the 2nd treatment, frontal cortex, hippocampus, amygdala and striatum were were dissected. The samples were stored at -80°C, and then lysed in Pierce RIPA sample buffer (Thermo Scientific, USA), supplemented with protease inhibitor cocktail (Roche Diagnostics, Germany) and boiled for 10 min. Equivalent amounts of protein extracts (20 mg) were analyzed by SDS–12% PAGE, followed by transfer of the proteins to polyvinylidene fluoride membrane. Blots were blocked in 5% fat-free milk in TBST buffer (Tris-Tween-buffered saline) and incubated in primary antibodies for one hour at room temperature. Primary antibodies included rabbit anti-GAP43 (ab75810, 1:2000; Abcam, UK), rabbit anti-PSD95 (ab238135, 1:2000; Abcam, UK), rabbit anti-Synaptophysin (ab32127, 1:2000; Abcam, UK), rabbit anti-SV2A (ab54351, 1:100; Abcam, UK) and mouse anti-β-Actin (8H10D10, 1:5000, Cell Signaling Technology). Blots were washed 3 times and incubated with horseradish peroxidase-conjugated secondary antibodies (1:5000, ABclonal, China) for 1 h, followed by repeated washing with TBST buffer. Proteins were visualized by using enhanced chemiluminescence (ChemiDoc Reader MP, Bio-Rad, USA). The amount of each phosphorylated protein was normalized to the amount of the corresponding total protein detected in the sample and measured by intensity of β-actin.

Representative immunoblots are shown in Supplementary Figures 1 and 2.

### f. Statistical analysis

The experimental data in all figures are expressed as the mean ± standard error of the mean (SEM). To determine intergroup differences, one-, two-, and three-way analyses of variance (ANOVA) were used as indicated. Bonferroni’s or Tukey’s multiple comparison tests were used to analyze post-hoc comparisons. p < 0.05 (two-tailed) was the criterion for significance. GraphPad Prism, version 10.2.3 software, was used for all statistical analyses. To assess the effect of treatment on each of the 4 synaptic protein levels across all 4 brain areas and on all 4 synaptic proteins within a specific brain area, nested analysis of variance (nested ANOVA) was used.

## 3. Results

### a. HTR

#### (i) D-Serine

PSIL (4.4mg/kg) induced a significant increase in HTR compared to vehicle control (p<0.0001, Fig 1A). DSER (3000 mg/kg) did not induce HTR. The combination of PSIL and DSER completely attenuated HTR (p<0.0001 vs. psilocybin), making it indifferent to VEH. To evaluate DSER at a lower dose, an experiment was conducted using PSIL (1.5mg/kg) and DSER (1500mg/kg). PSIL (1.5mg/kg) alone induced a significant increase in HTR compared to VEH (p<0.0001, Fig 1B). DSER (1500 mg/kg) alone did not induce an increase in HTR. The combination of PSIL (1.5mg/kg) and DSER (1500mg/kg) showed complete attenuation of HTR (p<0.0001, Fig 1B).

**Fig. 1:**
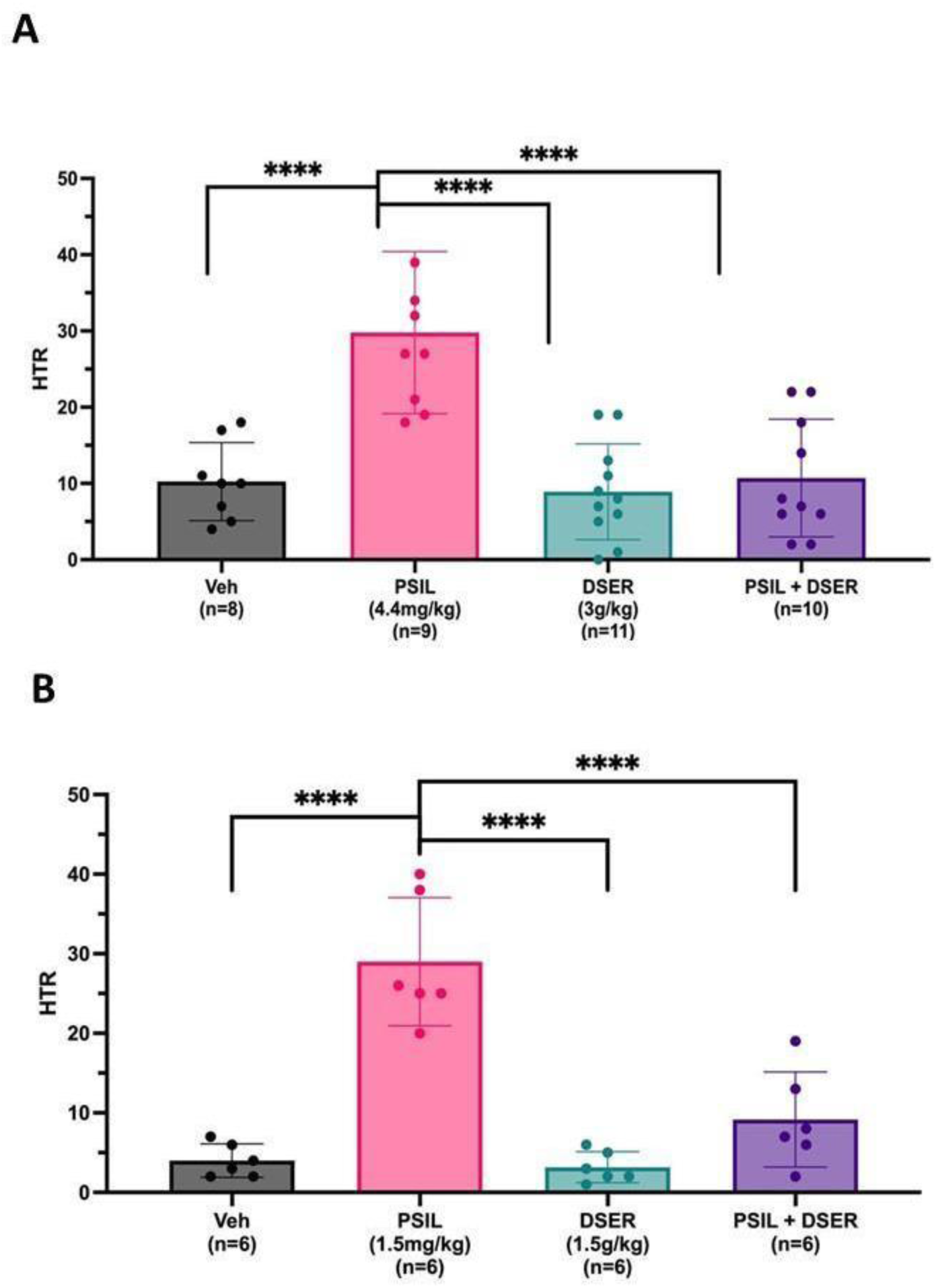
A. Effect of VEH, PSIL 4.4 mg/kg, DSER 3000 mg/kg, PSIL 4.4 mg/kg + DSER 3000 mg/kg on HTR over 30 minutes (F=15.33, df 3, 34, p<0.0001). B. Effect of VEH, PSIL 1.5 mg/kg, DSER 1500 mg/kg, PSIL 1.5 mg/kg + DSER 1500 mg/kg on HTR over 30 minutes (F=32.17, df 3, 20, p<0.0001). ****p<0.0001

#### (ii) D-Cycloserine

PSIL (4.4 mg/kg) induced a significant increase in HTR compared to saline control (p<0.0001). (Fig. 2A). DCS (320 mg/kg) did not affect HTR. The combination of PSIL and DCS attenuated HTR and induced a significantly lower number of head twitches compared to PSIL alone (p=0.0001), which was still significantly higher than the saline control (p=0.0023) (Fig. 2A). A dose-response experiment was performed to determine the lowest dose of DCS necessary to attenuate PSIL-induced HTR (Fig. 2B.) completely. DCS concentrations of 22.2mg/kg, 176.19mg/kg, and 440.47 mg/kg were used. With concentrations of 22.2mg/kg and 176.19mg/kg of DCS in combination with PSIL 4.4 mg/kg, there was no attenuation of HTR, and HTR levels were similar to those with PSIL alone. A combination of PSIL and DCS 440.47 mg/kg completely blocked HTR (p=<0.0001), and levels were similar to those of the saline control. It was decided to proceed with a DCS dose of 320 mg/kg in the following experiments.

**Fig. 2:**
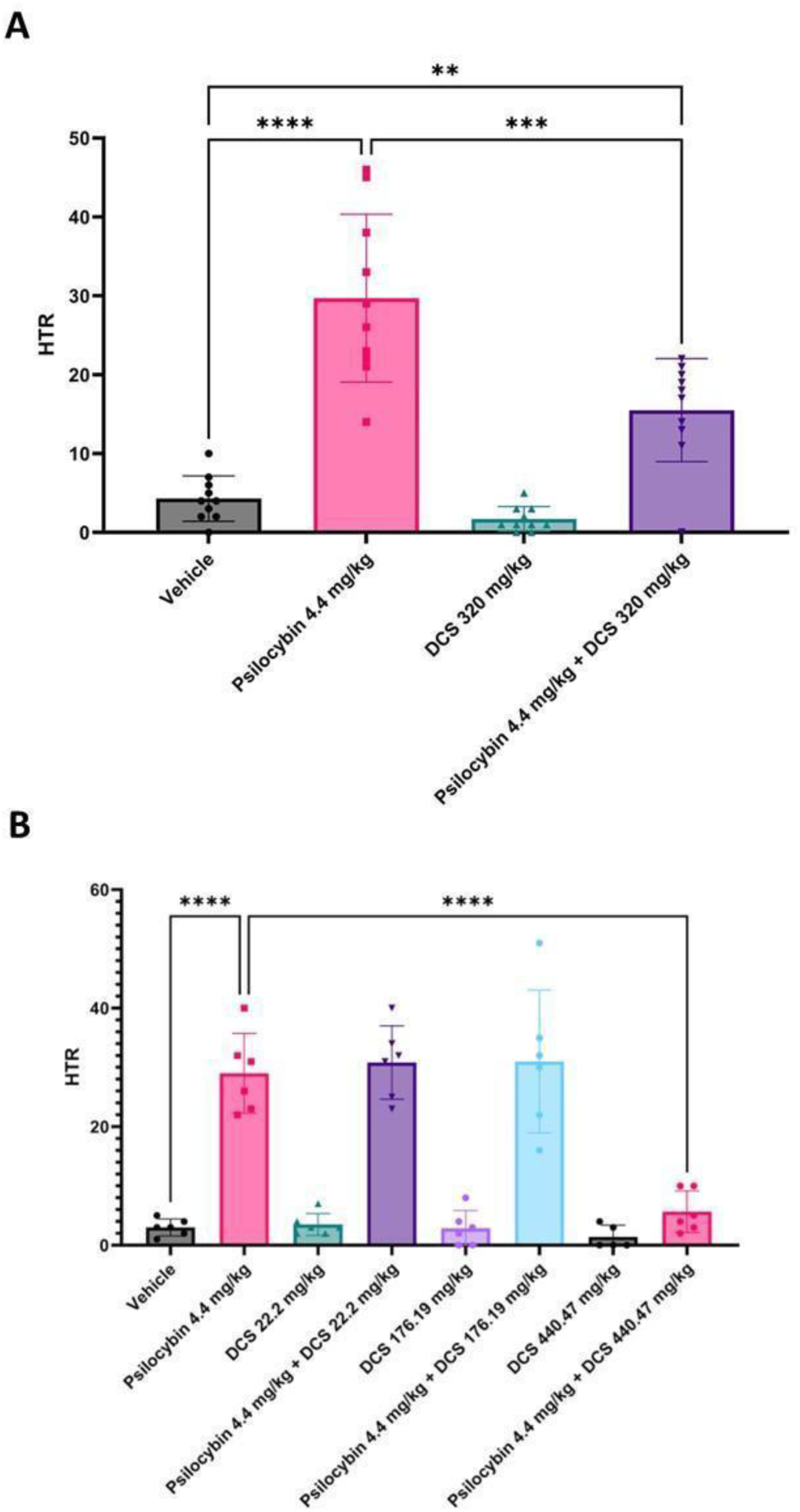
Effect of VEH, PSIL 4.4 mg/kg, DCS 320 mg/kg, PSIL 4.4 mg/kg + DCS 320 mg/kg on HTR over 20 minutes (F=39.10, df 3, 36, p<0.0001). B. Effect of VEH, PSIL 4.4 mg/kg, DCS 22.2 mg/kg, PSIL 4.4 mg/kg + DCS 22.2 mg/kg; DCS 176.19 mg/kg, PSIL 4.4 mg/kg + DCS 176.19 mg/kg; DCS 176.19 mg/kg, PSIL 4.4 mg/kg + DCS 176.19 mg/kg; on HTR over 20 minutes (F=35.04, df 7, 38, p<0.0001). **p<0.01; ***p<0.001; ****p<0.0001

### b. MK-801-induced hyperactivity

#### (i) D-Serine

Mice were injected intraperitoneally with VEH, PSIL 4.4 mg/kg, DSER 3000 mg/kg, or a combination of PSIL 4.4 mg/kg and DSER 3000 mg/kg 30 minutes before MK-801 injection. Locomotion was assessed in the open field for one hour, 30 minutes after MK-801 injection. DSER alone (3000 mg/kg) did not affect the hyperlocomotion induced by MK-801. Mice treated with PSIL+DSER prior to MK-801 had a statistically significant reduction in hyperlocomotion compared to MK+VEH, as seen in the distance traveled (p=0.0101, Fig. 3A).

**Fig. 3:**
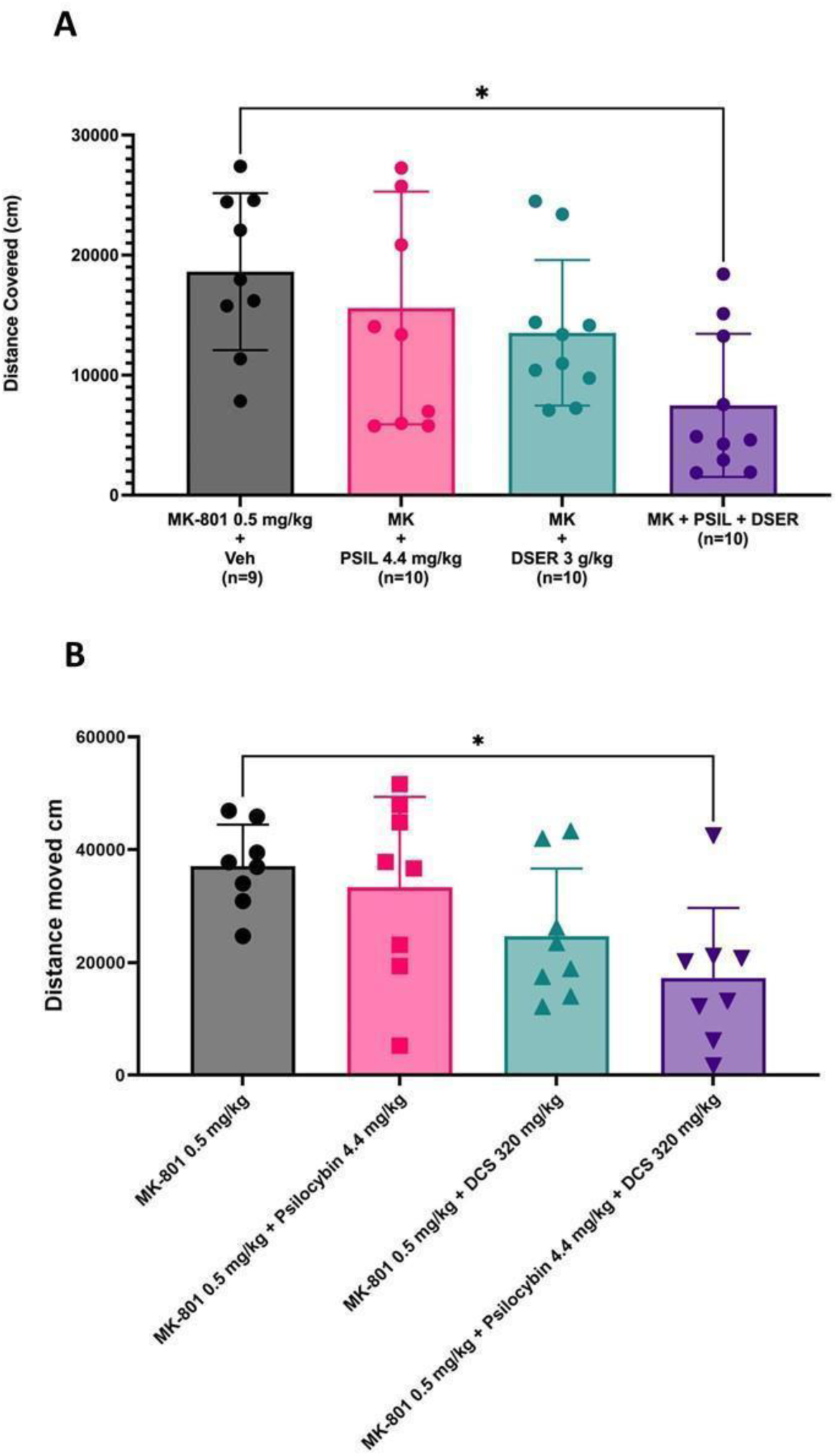
A. Effect of VEH, PSIL 4.4 mg/kg, DSER 3000 mg/kg, PSIL 4.4 mg/kg + DSER 3000 mg/kg on hyperlocomotion induced by MK-801 0.5 mg/kg (F=4.069, df 3, 35, p=0.0140). B. Effect of VEH, PSIL 4.4 mg/kg, DCS 320 mg/kg, PSIL 4.4 mg/kg + DCS 320 mg/kg on hyperlocomotion induced by MK-801 0.5 mg/kg (F=4.15, df 3, 28, p=0.02). *P<0.05.

#### (ii) D-Cycloserine

Mice were injected intraperitoneally with saline, psilocybin 4.4 mg/kg, DCS 320 mg/kg, or a combination of both. After 30 minutes, the mice were injected IP with MK 801 (0.5 mg/kg) and placed in the OFT arena for 60 minutes. DCS or PSIL alone did not significantly affect mobility induced by MK-801, but the combination of PSIL and DCS significantly reduced the MK-801-induced hyperactivity (p=0.0166) (Fig. 3B).

### c. Synaptic Proteins

#### (i) D-Serine

To assess the effects of PSIL and DSER, alone and in combination, on neuroplasticity, we assayed levels of 4 synaptic proteins (GAP43, PSD95, synaptophysin, and SV2A) in 4 brain areas (frontal cortex, hippocampus, amygdala, and striatum,) 11 days after a second treatment with VEH, PSIL 4.4 mg/kg, DSER 3000 mg/kg or PSIL 4.4 mg/kg + DSER 3000 mg/kg. In the frontal cortex, DSER induced a statistically significant increase in GAP43 levels compared to VEH (p=0.0068, Fig 4A). However, GAP43 levels following PSIL+DSER were similar to VEH (Fig. 4A). In the hippocampus, GAP43 levels induced by PSIL (p<0.01) and PSIL+DSER (p<0.01) were significantly higher than those induced by VEH (Fig. 4B). In the amygdala GAP43 levels induced by DSER alone (p<0.05) Iand PSIL+DSER (p<0.05) were significantly higher than VEH (Fig. 4C). Synaptophysin levels were significantly increased by DSER in the frontal cortex (p<0.05) and by PSIL in the hippocampus (p<0.05) (Supplementary Fig. 3). SV2A was increased by DSER (P<0.05) and by PSIL+DSER (p<0.05) in the hippocampus (Supplementary Fig. 4). PSD95 was not significantly altered by PSIL, DSER, or PSIL+DSER in any brain area (Supplementary Fig. 5) and no synaptic protein levels were significantly altered in the striatum (Fig. 4, Supplementary Figs. 3-5).

**Fig. 4:**
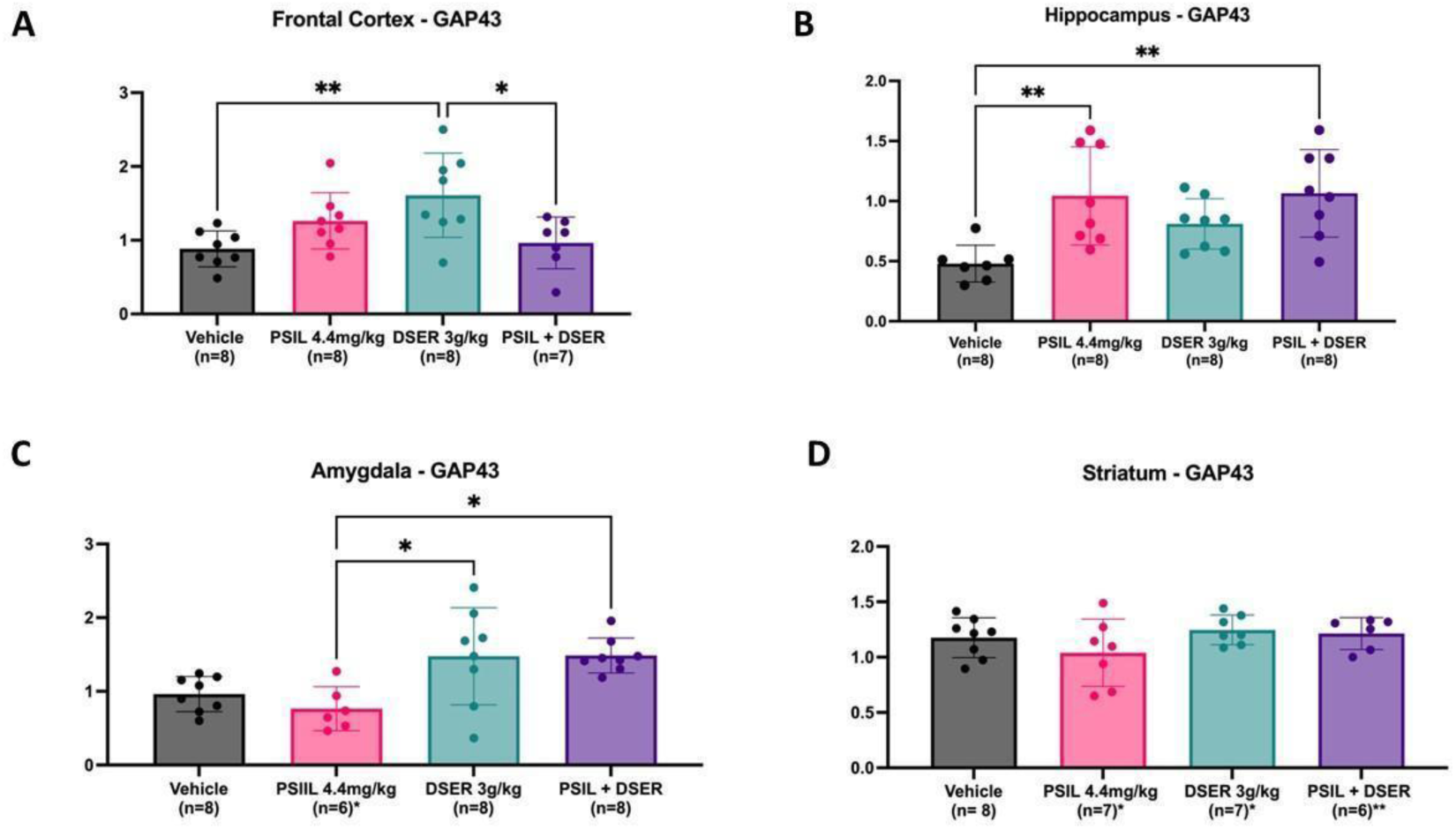
Effect of VEH, PSIL 4.4 mg/kg, DSER 3000 mg/kg, PSIL 4.4 mg/kg + DSER 3000 mg/kg on GAP43 levels in: A. Frontal Cortex (F=5.181, df 3, 27, p=0.0059), B. Hippocampus (F=5.775, df 3, 27, p=0.0035), C. Amygdala (F=5.795, df 3, 26, p=0.0036), D. Striatum (Not significant). *p<0.05; **p<0.01.

To assess the effect of the studied treatments on synaptic proteins across all 4 brain areas, we used nested ANOVA. For GAP43, there was a significant effect of DSER (p<0.001) and PSIL+DSER (p<0.01) (Fig. 5A and Supplementary Fig. 6); for synaptophysin of PSIL (p<0.001), DSER (p<0.01) and PSIL+DSER (p<0.05) (Supplementary Fig. 6). Performing nested ANOVA for all 4 synaptic proteins together within each brain area separately, a significant effect was observed in the hippocampus where the effect of PSIL+DSER to increase synaptic protein levels was significant (p<0.05) (Fig. 5, Supplementary Fig. 7) but not that of the other treatments. Treatment effects across synaptic proteins in other brain areas were not significant (Supplementary Fig. 7).

**Fig. 5:**
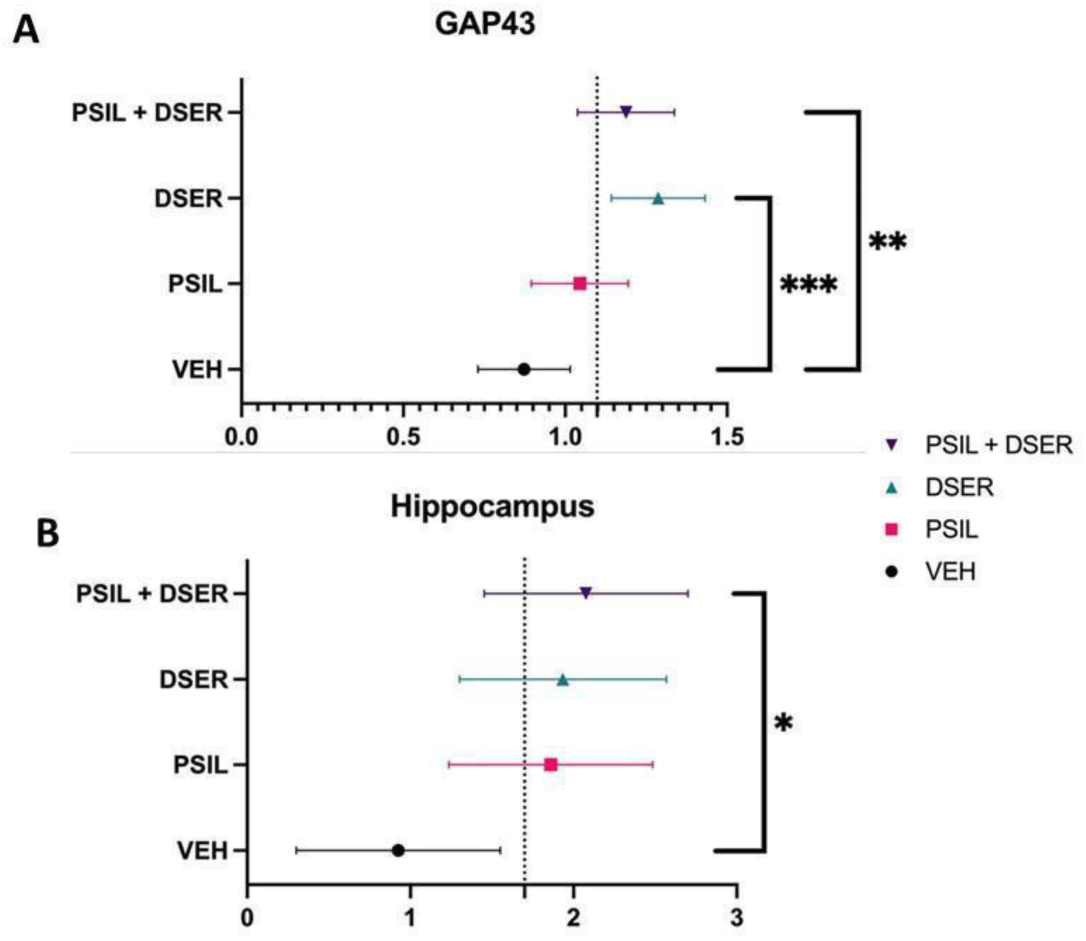
A. Nested ANOVA across 4 brain regions (frontal cortex, hippocampus, amygdala, striatum) showing effect of VEH, PSIL 4.4 mg/kg, DSER 3000 mg/kg, PSIL 4.4 mg/kg + DSER 3000 mg/kg. (F=6.126, df 3, 117, p=0.0007) on GAP43 levels. B. Nested ANOVA showing effect of VEH, PSIL 4.4 mg/kg, DSER 3000 mg/kg, PSIL 4.4 mg/kg + DSER 3000 mg/kg. on all 4 synaptic proteins (GAP43, PSD95, synaptophysin, SV2A) within the hippocampus (F=2.747, df 3, 123, p=0.0458). *p<0.05; **p<0.01, ***p<0.001

#### (ii) D-Cycloserine

Western blot analysis of the synaptic proteins GAP43, PSD95, synaptophysin and SV2A was performed on frontal cortex, amygdala, hippocampus, and striatum samples from mice injected with VEH, PSIL 4.4 mg/kg, DCS 320 mg/kg, or a combination of PSIL and DCS 12 days after treatment. There was a significant increase in PSD95 in the frontal cortex of mice that received a combination of PSIL and DCS (p=0.03) (Fig. 6A) but not in the other brain areas (Supplementary Fig. 8). When PSD65 levels were evaluated across all 4 brain areas by nested ANOVA, the results showed a significant effect of PSIL+DCS to increase PSD95 levels (Fig. 6B). There were no significant effects of any treatments on GAP43, synaptophysin or SV2A levels in any of the 4 brain regions (Supplementary Figs. 8-9). Nested ANOVA did not reveal a significant effect of the treatments administered on GAP43, synaptophysin or SV2A across the 4 brain regions (Supplementary Fig. 10) and not on all 4 synaptic proteins within any of the 4 brain regions (Supplementary Fig. 11).

**Fig. 6.:**
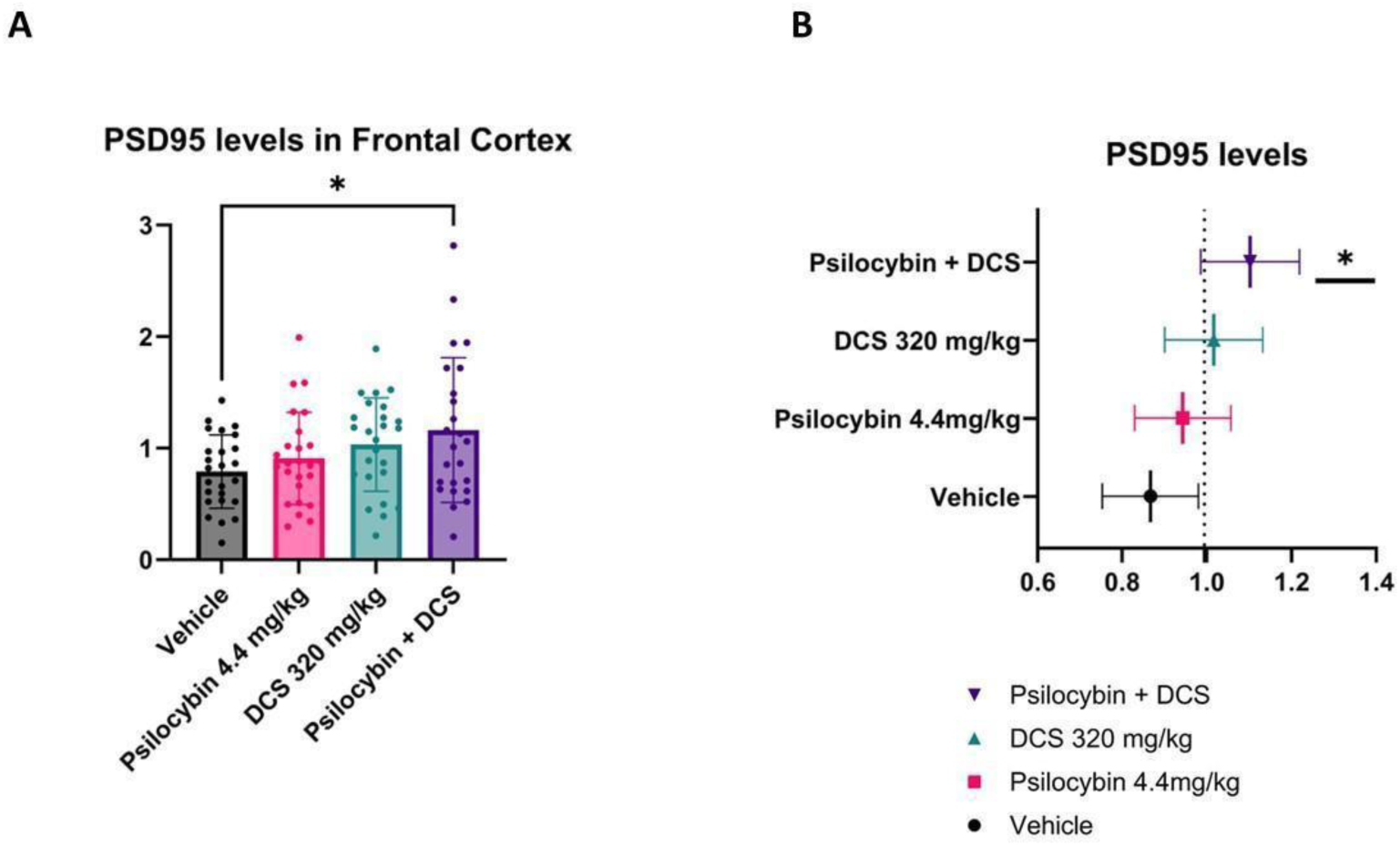
A. PSD95 levels in the frontal cortex 11 days after treatment with VEH, PSIL 4.4 mg/kg. DCS 320 mg/kg, PSIL 4.4 mg/kg + DCS 320 mg/kg (F=2.30, df 3, 95, p=0.04). B. Nested ANOVA of PSD95 levels across 4 brain regions (Frontal cortex, hippocampus, amygdala, striatum) (F=3.02, df 3, 95, p=0.03). *p<0.05

## 4. Discussion

To the best of our knowledge, these are the first experiments performed to assess the effects of concomitant administration of an SP and an NMDAR modulator. Overall, the present proof-of-concept findings indicate that both DSER and DCS have the potential to improve safety and efficacy parameters when administered concurrently with psilocybin.

As expected, HTR, which is an animal proxy of the psychedelic trip in humans ^56^, was significantly elicited by psilocybin. DSER and DCS alone did not affect HTR. Furthermore, both DSER and DCS, when administrated in conjunction with psilocybin, dose-dependently reduced HTR up to a complete block with DSER 3g/kg and DCS 440.47mg/kg doses. These findings are in line with previous work in murine models. The administration of the NMDAR antagonist phencyclidine induces HTR ^57^. Both competitive and noncompetitive NMDAR antagonists (MK-801, ketamine, dextromethorphan) markedly enhance serotonin-induced HTR, while N-methyl-D-aspartate (NMDA) itself inhibits HTR generation ^58^. Our results indicate that NMDAR-GMS modulator administration may potentially be used to calibrate or even abolish the hallucinogenic effects registered during psilocybin treatment.

In contrast to HTR generation, we found that psilocybin did not significantly exacerbate (but neither alleviated) MK-801-induced hyperlocomotion. In contrast, and unlike each of the compounds given separately, combined psilocybin + DSER, as well as psilocybin + DCS administration, significantly reduced the hyperlocomotion induced by MK-801. Since this type of hyperactivity is widely considered a reliable animal model of psychotic symptoms ^53, 54, 59^, our findings suggest that psilocybin may be employed in psychotic disorders without carrying an *a priori* major risk for psychotic exacerbation. Additional support for this concept may be drawn from naturalistic studies involving SP administration to autism and schizophrenia patients performed during the 1950s and 60s ^60–62^. Moreover, while it has been proposed that SP may alleviate negative symptoms and cognitive deficits of schizophrenia ^3^, the present results indicate that the combined psilocybin-NMDAR modulator treatment may also achieve improvements in psychosis and positive symptoms. Taken together with the findings on HTR, the results of the present study confirm the hypothesis that the combined treatment may overall offer a protective effect against undesirable hallucinatory and pro-psychotic side effects that may be generated by psilocybin.

The main potentially therapeutic mechanisms by which SP and NMDAR modulators may exert synergistic effects are the amplification and diversification of neuroplastic effects ^11^. Cardinal common denominators of these processes are the activation of glutamatergic ionotropic receptors and synaptogenesis. In the present study, in addition to testing the immediate behavioral effects of psilocybin, DSER, DCS, and their combinations, prolonged effects on neuroplasticity were tested by performing western blot analysis in brain samples for the synaptic proteins GAP43, PSD95, synaptophysin, and SV2A. Due to technical limitations, the methodologies employed differed slightly; for DSER-related experiments, the samples were obtained 11 days after a second treatment administration, while in the DCS experiments, the samples were obtained 12 days after a single administration of the compounds. The observed results were heterogeneous, with psilocybin, DSER, and DCS affecting the levels of some of the measured proteins in some of the brain areas examined. Overall, for the psilocybin-DSER combination, significant synergistic effects were found across all brain regions examined for GAP43 levels and for all the proteins measured in the hippocampus. Combined psilocybin-DCS administration resulted in a significant increase in PSD95 in all brain areas examined. Overall, while identifying a signal for synergistic effects of psilocybin and DSER/DCS, the present results may have been influenced by a number of factors, including dosages and methodological issues thus warranting further research.

PSD95 is known as a major regulator of synaptic maturation by interacting, stabilizing, and trafficking NMDAR to the postsynaptic membrane ^63, 64^ and may thus be used as a synaptogenesis marker. Moreover, PSD95 disruption and insufficiency have recently been associated with cognitive and learning deficits in schizophrenia ^65^. The evidence that combined treatment with psilocybin and DCS leads to up-regulation of PSD95 levels in the brain, in addition to blocking the hyperlocomotion caused by MK-801, suggests a therapeutic potential of the proposed treatment in schizophrenia.

While DSER and DCS may defend against SP-induced undesirable effects, the issue of their own side effects and dosage should be considered. DSER has an excellent safety profile. The only significant concern seems to be DSER-induced nephrotoxicity that has been reported in rats but not in any other species. Even in rats, DSER-related tubular necrosis is dose dependent and reversible; and does not appear to be present at doses producing an acute Cmax of <2000 nmol/ml ^66, 67^. In contrast, the acute Cmax of DSER 120 mg/kg, the highest dose tested in humans, is ∼500 nmol/ml ^67^. Across all published DSER human studies, including ∼500 DSER receivers, only one person has been reported to have abnormal renal values that fully resolved within a few days of stopping treatment ^68^. Nevertheless, a significant challenge for DSER use is low oral bioavailability. On oral administration, DSER is substantially catabolized by D-amino acid oxidase ^31, 69^, diminishing its bioavailability and necessitating, as seen also in the present study, the administration of gram-level doses. In view of this limitation, the ideal dosage and mode of administration of DSER may involve the synchronous administration of compounds with overlapping mechanisms of action.

DCS actions at NMDAR are particularly complex since its effects on NMDAR channel opening vary by dose and NMDAR subtype. DCS is a weak partial agonist at GluN2A- and GluN2B-containing NMDAR, with greater agonist efficacy at GluN2C and GluN2D-containing receptors ^70, 71^. The enhanced plasticity reported with DCS (35-40) and observed with high doses also in our study may involve changes in subunit composition as well as changes in expression of molecules involved in intracellular pathways that mediate plasticity and stimulation of neurogenesis ^72, 73^. It has been suggested that DCS may preferentially facilitate or inhibit NMDAR subtypes that may have specific clinical relevance ^48, 72, 73^ and that high dose DCS may attenuate executive cognitive deficits associated with suicide risk ^48^. The low specificity for different NR2 subunits and tachyphylaxis are, in fact, the main DCS limitations. Newer approaches should explore the efficacy of dual treatment combining DCS with other treatments, such as neuropeptides ^74^, transcranial magnetic stimulation ^75^ and, in our proposition-SP ^11^.

In regard to DCS side effects, although associated in early chronic tuberculosis treatment-related reports with detrimental neuropsychiatric effects, no propensity for dissociative phenomena or suicidality ^47–49^, abuse ^76^, or neurotoxicity ^77^ that have been reported with NMDAR direct channel antagonists such as ketamine, have been associated with DCS. Moreover, it was shown that the rate of spontaneous mutations conferring resistance to DCS (mutation rate) is ultra-low in *M. tuberculosis* (ca. 10^-11^) ^78^. In fact, DCS has been used for six decades without significant appearance and dissemination of antibiotic resistant strains, making it an ideal model compound to understand what drives resistance evasion ^78^.

The present study has a number of limitations. The study aimed of detecting safety and efficacy signals of the combined psilocybin-DSER/DCS treatments as compared to the effects of the single components. However, the single acute administration paradigm employed in the present study may not be the optimal form of administration of the combined treatment. Acute administration of psilocybin, DSER and DCS were all shown to elicit neuroplastic effects in animal models, while in clinical research single administration of psilocybin and daily administration of DSER or DCS for a number of weeks have been mainly assessed. Acute, intermittent and prolonged treatment, eventually using reduced dose strategies, are all forms of administration that may be relevant to the proposed combined treatments ^3, 11^. Further animal work exploring additional administration paradigms is warranted. Ultimately, the dose and frequency of administration will likely be critical to maximize efficacy while minimizing undesirable side effects. Additional drawbacks of the present study include the relatively small animal samples and the lack of assessment of additional relevant markers, e. g. GLU and BDNF levels.

Overall, our results suggest that the combined SP-NMDAR modulator administration could represent an innovative synergistic treatment approach that may hold efficacy and safety advantages versus treatment with SP or DSER/DCS alone. Randomized controlled clinical trials are the key test to our hypothesis. If confirmed, the present results indicate that SP treatment could be optimized, leading to its widespread utilization across a wider array of neuropsychiatric disorders.

## Supporting information

Ben Tal - Supplemental Information

## Acknowledgments

The authors acknowledge the expert technical assistance of Masha Chaykin, the assistance of May Ben Ari, Miles Guralnik, Ethan Hamid, Bar Orian, and Chloe Shevakh with experiments, and the valuable advice of Orr Shahar and Gilly Wolf.

## Support

This work was supported in part by Negev Labs

## Conflict of Interest

PCT WO 2024/052895 A1 (Inventors: BL, UHL), submitted by Hadasit Medical Research and Development Ltd and Ezrath Nashim - the Rabanit Herzog Memorial Hospital discloses SP-NMDAR modulator combination therapy for psychiatric and other disorders.

### Author Contributions

Conception and design: UHL, BL Acquisition of data: TBT, IP, AB Analysis and interpretation of data: UHL, BL, TBT, IP, AB

Drafting the article or revising it critically for important intellectual content: UHL, BL, TBT, IP, AB Final approval of the version to be published: All authors

### Data Availability

Data will be provided for further analysis to qualified investigators. Please contact

