## Supplementary material for "Synergistic Behavioral and Neuroplastic Effects of Psilocybin-NMDAR Modulator Administration": Ben Tal - Supplemental Information

**Ben Tal et al.:**

### SUPPLEMENTARY INFORMATION

**Supplementary Fig. 1**

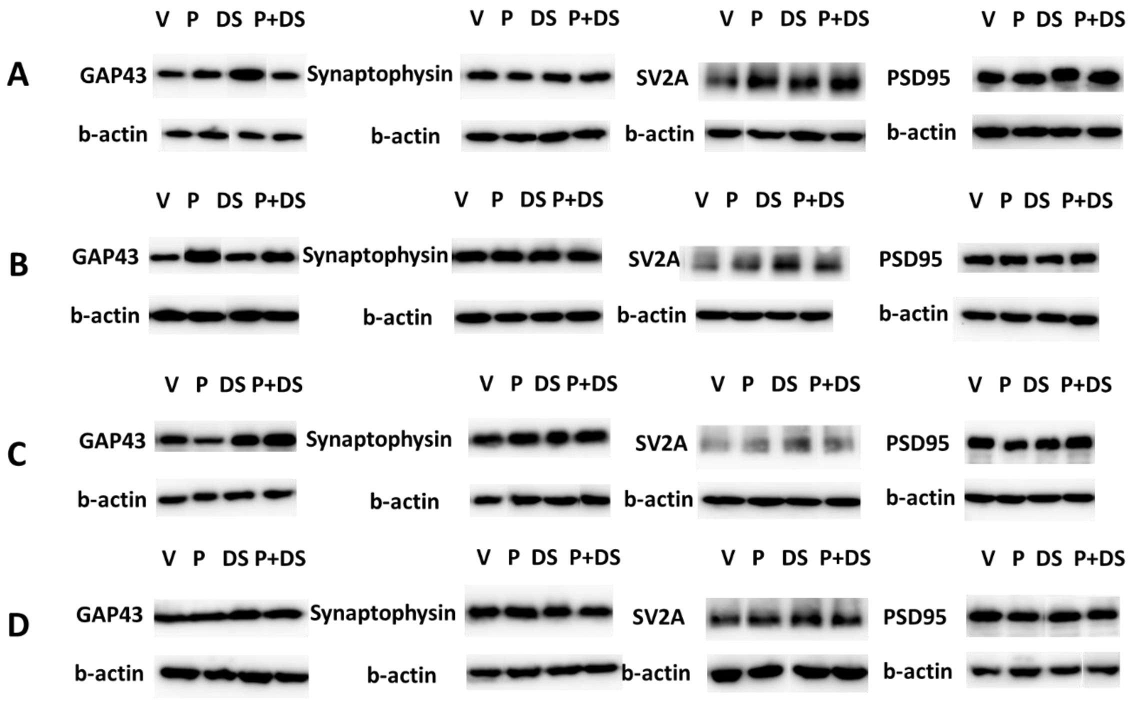


**Supplementary Fig. 1.** Representative immunoblots of GAP43, Synaptophysin, SV2A, PSD95 and b-actin protein expression in the frontal cortex (A), hippocampus (B), amygdala (C) and striatum (D), after V (Vehicle), P (Psilocybin), DS (DSER) and P+DS (Psilocybin and DSER) treatments.

**Supplementary Fig. 2**

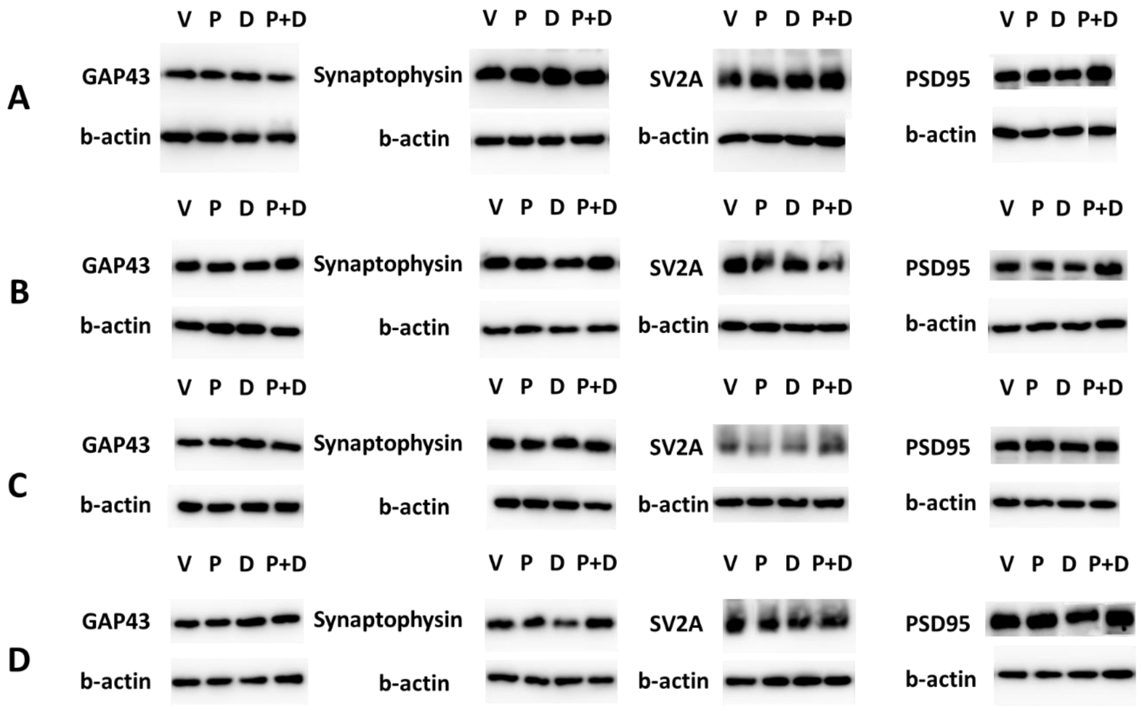


**Supplementary Fig. 2.** Representative immunoblots of GAP43, Synaptophysin, SV2A, PSD95 and b-actin protein expression in the frontal cortex (A), hippocampus (B), amygdala (C) and striatum (D), after V (Vehicle), P (Psilocybin), D (DCS) and P+D (Psilocybin and DCS) treatments.

**Supplementary Fig. 3**

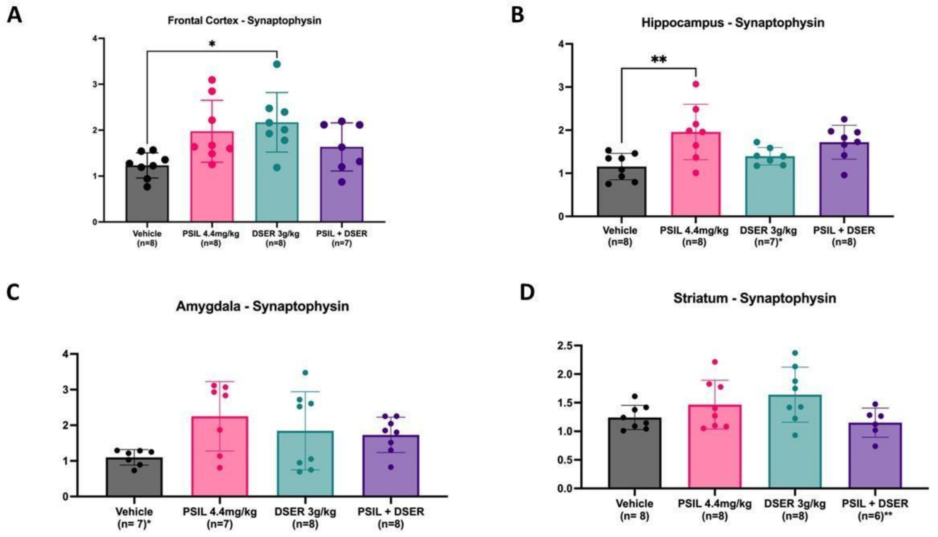


**Supplementary Fig. 3:** Effect of VEH, PSIL 4.4 mg/kg, DSER 3000 mg/kg, PSIL 4.4 mg/kg + DSER 3000 mg/kg on synaptophysin levels in: A. Frontal Cortex (F=4.399, df 3, 27, p=0.0121), B. Hippocampus (F=5.482, df 3, 27, p=0.0045), C. Amygdala (Not significant), D. Striatum (Not significant).**p<0.01.

**Supplementary Fig. 4**

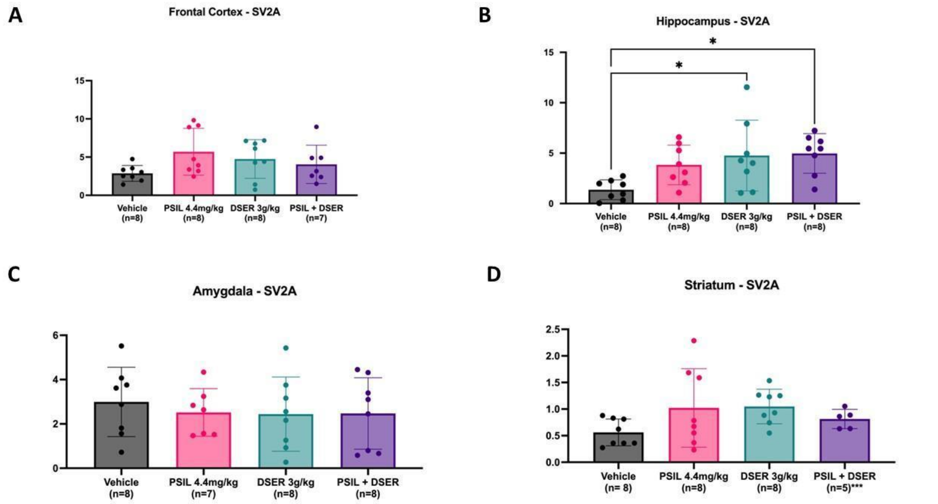


**Supplementary Fig. 4:** Effect of VEH, PSIL 4.4 mg/kg, DSER 3000 mg/kg, PSIL 4.4 mg/kg + DSER 3000 mg/kg on SV2A levels in: A. Frontal Cortex, B. Hippocampus (F=4.148, df 3, 28, p=0.0150), C. Amygdala , D. Striatum. No other differences are statistically significant. *p<0.05

**Supplementary Fig. 5**

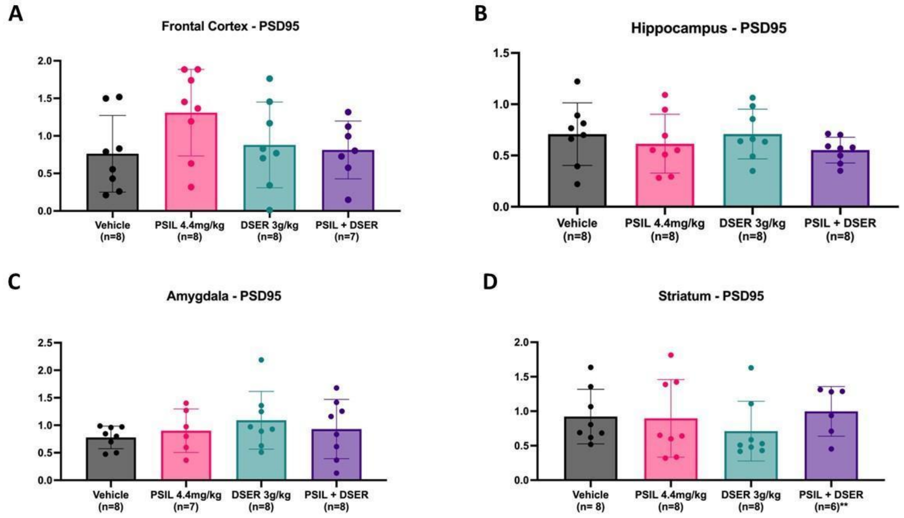


**Supplementary Fig. 5:** Effect of VEH, PSIL 4.4 mg/kg, DSER 3000 mg/kg, PSIL 4.4 mg/kg + DSER 3000 mg/kg on PSD95 levels in: A. Frontal Cortex, B. Hippocampus, C. Amygdala, D. Striatum. (No differences statistically significant).

**Supplementary Fig. 6**

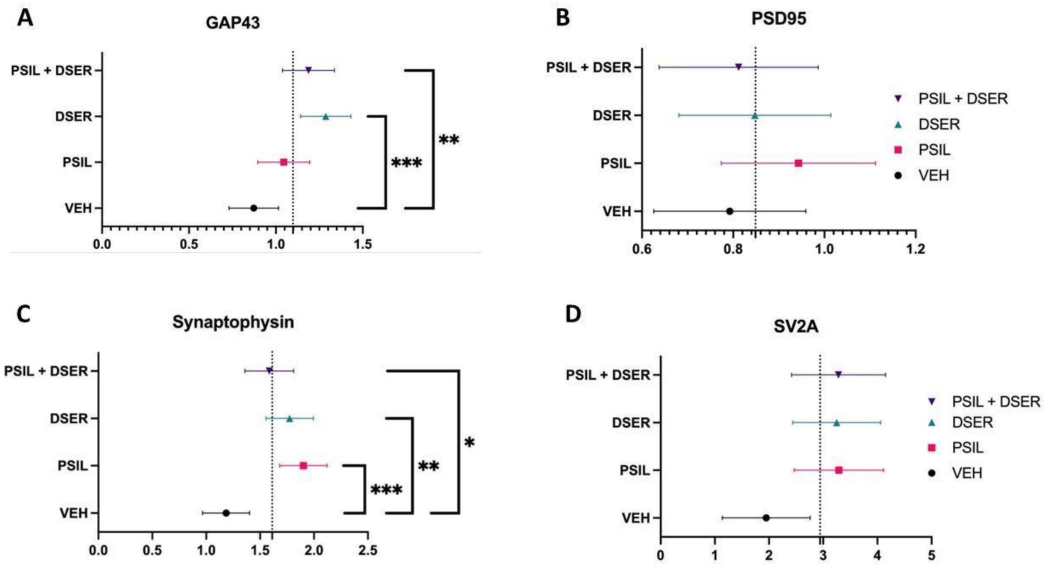


**Supplementary Fig. 6:** Nested ANOVA across 4 brain areas (frontal cortex, hippocampus), amygdala, striatum) of the synaptic proteins, GAP43 (F=6.126, df 3, 117, p=0.0007), PSD95 (Not significant), synaptophysin (F=8.554, df 3, 28, p=0.0003), SV2A (Not significant) 11 days after treatment with VEH, PSIL 4.4 mg/kg, DSER 3000 mg/kg, PSIL 4.4 mg/kg + DSER 3000 mg/kg. *p<0.05, **p<0.01; ***p<0.001

**Supplementary Fig. 7**

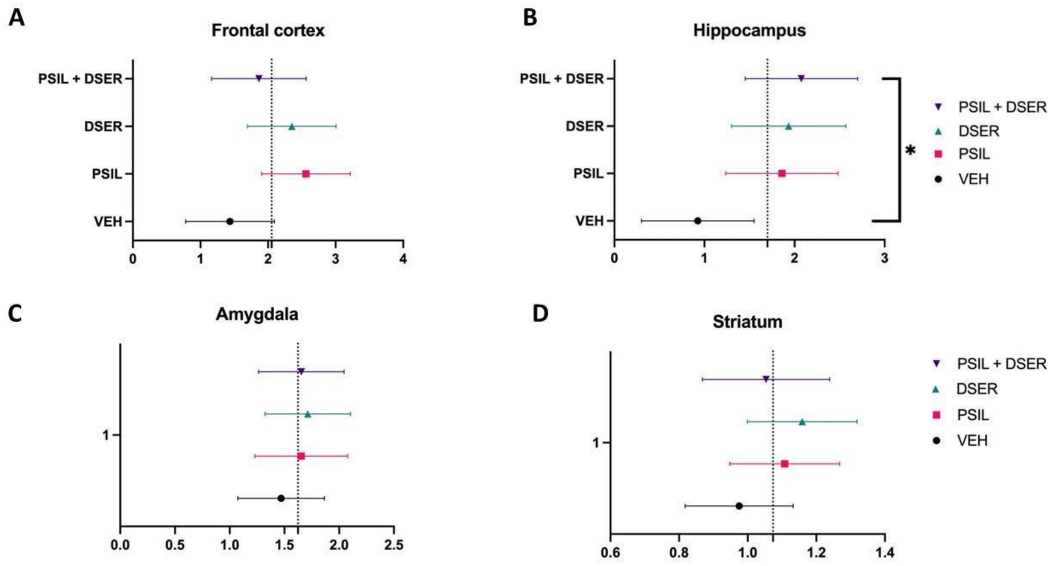


**Supplementary Fig. 7:** Nested ANOVA across the synaptic proteins GAP43, PSD95, synaptophysin and SV2A in A. frontal cortex (Not significant), B. hippocampus (F=2.747, df 3, 123, p=0.0458), amygdala (Not significant) and striatum (Not significant) 11 days after treatment with VEH, PSIL 4.4 mg/kg, DSER 3000 mg/kg, PSIL 4.4 mg/kg + DSER 3000 mg/kg. *p<0.05.

**Supplementary Fig. 8**

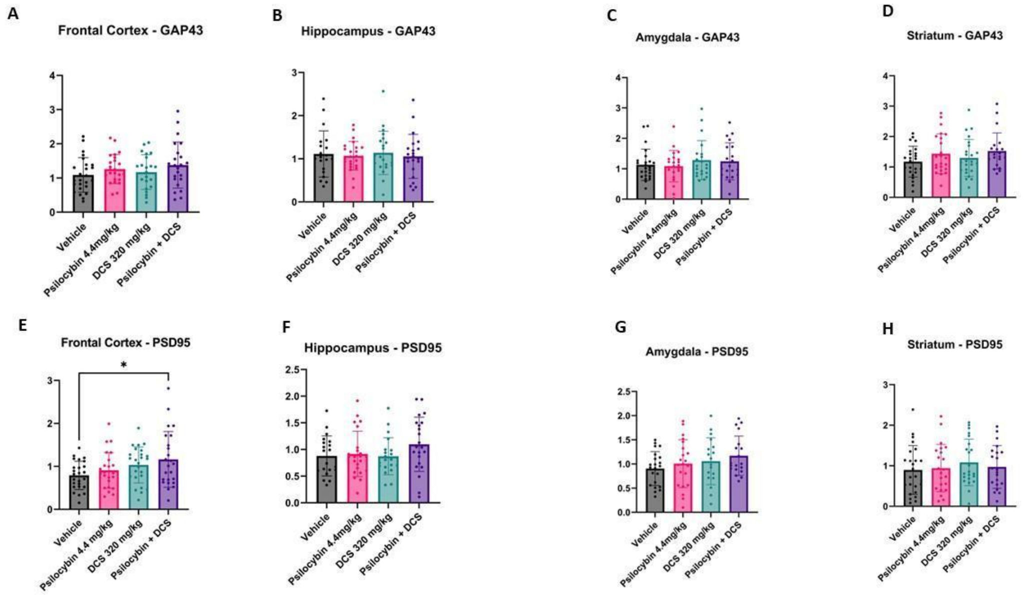


**Supplementary Fig. 8:** Effect of VEH, PSIL 4.4 mg/kg, DCS 320 mg/kg, PSIL 4.4 mg/kg + DCS 320 mg/kg on A. GAP43 levels in: A. Frontal Cortex, B. Hippocampus, C. Amygdala, D. Striatum; and on PSD95 levels in: E. Frontal Cortex (F=2.90, df 3, 95, p=0.04), F. Hippocampus, G. Amygdala, H. Striatum; (No other differences statistically significant). *p<0.05

### Supplementary Fig. 9

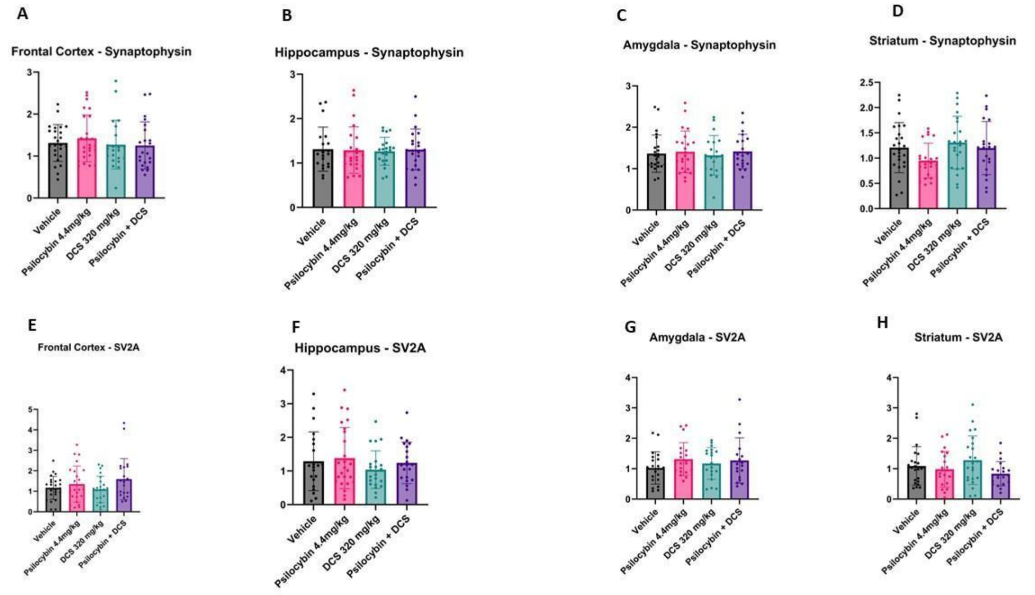


**Supplementary Fig. 9:** Effect of VEH, PSIL 4.4 mg/kg, DCS 320 mg/kg, PSIL 4.4 mg/kg + DCS 320 mg/kg on A. synaptophysin levels in: A. Frontal Cortex, B. Hippocampus, C. Amygdala, D. Striatum; and on SV2A levels in: E. Frontal Cortex, F. Hippocampus, G. Amygdala, H. Striatum; (No differences statistically significant).

**Supplementary Fig. 10**

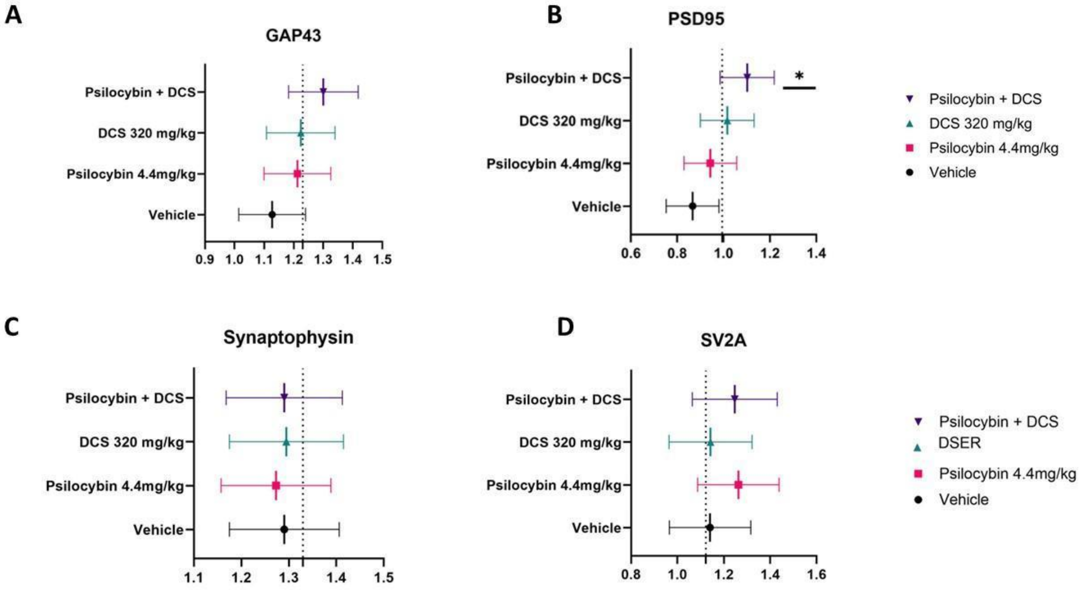


**Supplementary Fig. 10:** Nested ANOVA of the synaptic proteins GAP43, PSD95 (F=3.02, df 3, 95, p=0.03), synaptophysin across 4 brain regions (frontal cortex, hippocampus, amygdala and striatum) 11 days after treatment with VEH, PSIL 4.4 mg/kg, DCS 320 mg/kg, PSIL 4.4 mg/kg + DCS 320 mg/kg. *p<0.05.

**Supplementary Fig. 11**

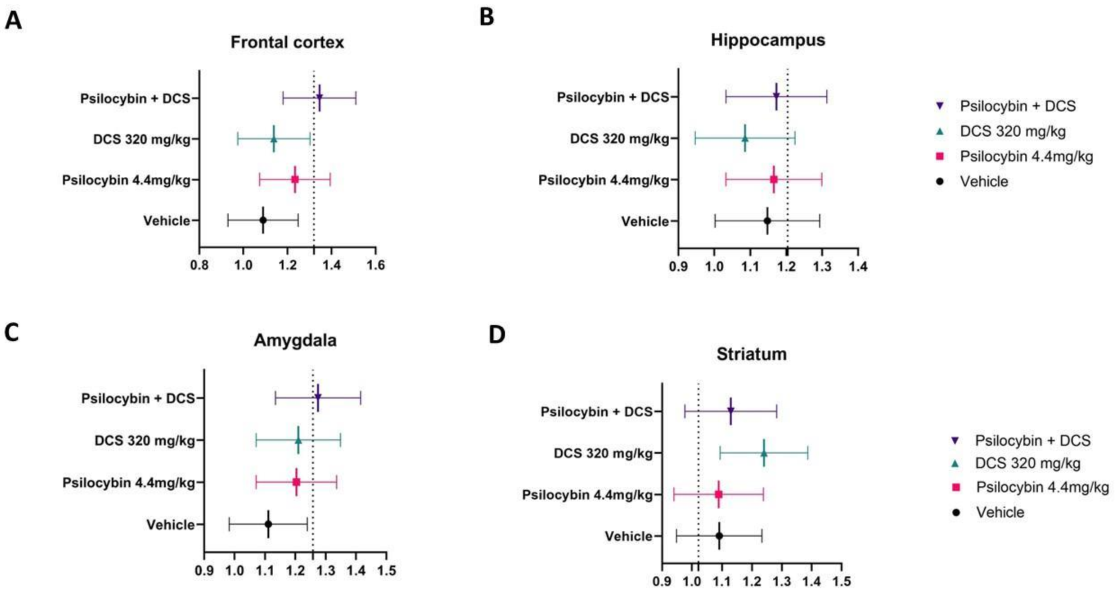


**Supplementary Fig. 11:** Nested ANOVA across the synaptic proteins GAP43, PSD95, synaptophysin and SV2A in frontal cortex, hippocampus, amygdala and striatum 12 days after treatment with VEH, PSIL 4.4 mg/kg, DCS 320 mg/kg, PSIL 4.4 mg/kg + DCS 320 mg/kg. (No differences statistically significant).
